# Sarcomeric Remodelling in Human Heart Failure unraveled by single molecule long read sequencing

**DOI:** 10.1101/2025.02.28.640805

**Authors:** Jan Haas, Sarah Schudy, Benedikt Rauscher, Ana Munoz, Steffen Roßkopf, Christoph Reich, Gizem Donmez Yalcin, Abdullah Yalcin, Timon Seeger, Manuel H. Taft, Marc Freichel, Dirk Grimm, Dietmar Manstein, Johannes Backs, Norbert Frey, Lars Steinmetz, Benjamin Meder

## Abstract

Dysregulation of alternative splicing – mediated by factors such as RBM20 or SLM2 – can affect proper gene isoform control disrupts gene isoform homeostasis and underpins severe cardiomyopathy in both animal models and patients. Although innovative therapies target various sarcomeric components, the impact of isoform switching in cardiac disease remains poorly understood. Here, we applied nanopore long-read sequencing to map the full-length transcriptome of left ventricular tissue from thirteen nonfailing controls, ten patients with dilated cardiomyopathy (DCM), and ten with ischemic cardiomyopathy (ICM). Our analysis identified 78,520 transcripts, 31% of which represent novel isoforms of known genes. Notably, the transcriptomes of DCM and ICM were largely indistinguishable, indicating that end-stage heart failure is characterized by a convergent isoform landscape, irrespective of disease etiology. Among 11 prototypical sarcomere genes, 10 displayed highly significant isoform shifts (p= 5.23×10^−45^ – 2.89×10^−200^). Focusing on tropomyosin, we observed that while the predominant cardiac gene *TPM1* showed moderate up-regulation of its transcript isoforms, transcripts derived from *TPM3*—typically expressed at lower levels in the healthy heart—were markedly increased in heart failure.

## Introduction

Heart failure is a severe medical condition in which the heart is weakened and unable to pump blood properly (Ponikowski *et al*, 2016). It is caused by a variety of diseases: e.g. arrhythmia, toxic damage, ischemic heart disease, and cardiomyopathies. Cardiomyopathy is defined as heart muscle disease with structural and/or electrical dysfunction with four main subgroups, with dilated cardiomyopathy (DCM) being the most prevalent one (Maron *et al*, 2006) (Elliott, 2023). Besides secondary causes, 30-50% of the DCM cases are known to be genetic (Hershberger *et al*, 2013) involving mostly sarcomeric and z-disc associated genes (McNally & Mestroni, 2017) (Millat *et al*, 2011) (van Spaendonck-Zwarts *et al*, 2013) (Hershberger *et al*, 2018). Ischemic cardiomyopathy (ICM) derives from myocardial infarction or coronary artery disease (Dang *et al*, 2020). Underlying causes also involve environmental factors and genetics.

Several cases are known in which aberrant alternative splicing contributes to cardiomyopathies (Kong *et al*, 2010) (Sweet *et al*, 2018). Splicing factors such as RNA-binding motif protein 20 (RBM20) determine the physiological mRNA landscape formation, and rare variants in the *RBM20* gene explain up to 6% of genetic DCM cases (Koelemen *et al*, 2021). Putative splice regulators with relevance during heart failure include Sam68-like mammalian protein 2 (SLM2) or RBFox 1. In the human heart, SLM2 binds to important transcripts of sarcomere constituents, such as those encoding myosin light chain 2 (MYL2), troponin I3 (TNNI3), troponin T2 (TNNT2), tropomyosin 1/2 (TPM1/2), and titin (TTN) (Boeckel *et al*, 2022). Due to the complex structure and variety in the length of sarcomeric genes, the complete coverage of the exon junctions is challenging (Uapinyoying *et al*, 2020), leading to increased efforts to develop novel techniques to identify splicing events in a more accurate manner. Nanopore long-read sequencing is a rather new technique known as third-generation sequencing, enabling researchers to sequence extremely long stretches of nucleic acids on at single-nucleotide resolution (Wu *et al*, 2023).

The technology leverages nanoscale protein pores embedded in an electrically resistant polymer membrane, which act as biosensors to permit the passage of a single nucleic acid strand at a time. This enables the sequencing of extended read lengths, facilitating the differentiation of highly similar isoforms, such as tropomyosins.

Tropomyosins are actin-binding proteins in the sarcomere and are encoded by over 40 different isoforms (Li *et al*, 2011). In muscle cells, tropomyosin contributes to the regulation of sarcomere contraction by sliding along the actin filament in response to calcium (Ca^2+^) binding to the troponin complex (composed of troponin C, T, I encoded by *TNNC1*, *TNNT2*, *TNNI3* respectively). This movement exposes the myosin-binding sites on the actin filament, enabling muscle contraction (Craig & Lehman, 2001). The functional isoforms of tropomyosins in the heart are not well defined and differ between studies and authors. Several reports have highlighted the varying contributions of *TPM1* gene products to heart disease, with multiple variants identified as causative factors (Walsh *et al*, 2017). During rat heart development, full-length isoforms of *TPM1* were identified using nanopore long-read sequencing, with specific exon expression patterns associated with distinct developmental stages (Cao *et al*, 2021). Functionally, it is known that tropomyosin isoforms impact on the actin-myosin binding by altering the maximum speed of the myosin motors or their properties of calcium sensitivity (Nefedova *et al*, 2022) (Farman *et al*, 2018; Pertici *et al*, 2021; Reindl *et al*, 2022).

In this study, we delineate for the first time the isoform landscape of different forms of heart failure by single molecule sequencing. We show the remarkable changes in isoform expression, especially remodelling the sarcomere with its core components. Functionally, we show that different isoforms of tropomyosin have altered calcium sensitivity, reflecting their potential adaptation to the changes of cellular requirements.

## Results

### Gene expression analysis in heart failure patients

In order to unravel the transcriptomic landscape in heart failure patients, we have performed cDNA full length sequencing for 13 controls, 10 DCM patients, and 10 ICM patients. Full-length transcriptome analysis was done using FLAMES (https://github.com/LuyiTian/FLAMES). During differential gene expression analysis (DESeq2), we established a stringent significance threshold through p-value permutation (100 iterations). When samples from heart failure patients (DCM and ICM) were compared to controls, out of 22,622 genes, 872 genes were significantly up-regulated, while 2,224 genes were significantly down-regulated based on the calculated p-value cut-off of 3.2×10^−5^ (**Fig 1A**). Based on significance, the most downregulated genes in heart failure (HF) patients were *PDIA5* (padj=6×10^−39^)*, ADAMTS4* (padj=9×10^−28^) *and MMRN2* (padj=6×10^−27^). The most upregulated genes were shown to be *PCCB* (padj=5×10^−26^)*, MRPL44* (padj=1×10^−23^) and *DLAT* (padj=1×10^−23^). The largest fold-change was associated with an increase in gene expression of *CA3* (log2FoldChange=3.16) and a decrease in gene expression of *S100A9* (log2FoldChange=-4.03) in HF patients (**Fig. 1A**). *NPPA*, prototypical heart failure biomarker, was also strongly upregulated on its isoform level (padj=3×10^−6^; log2FoldChange=2.5). When comparing DCM and ICM, no differentially expressed genes were observed (p-value cutoff = 2.8×10^−5^) (**Fig. 1B**).

**Figure 1:**
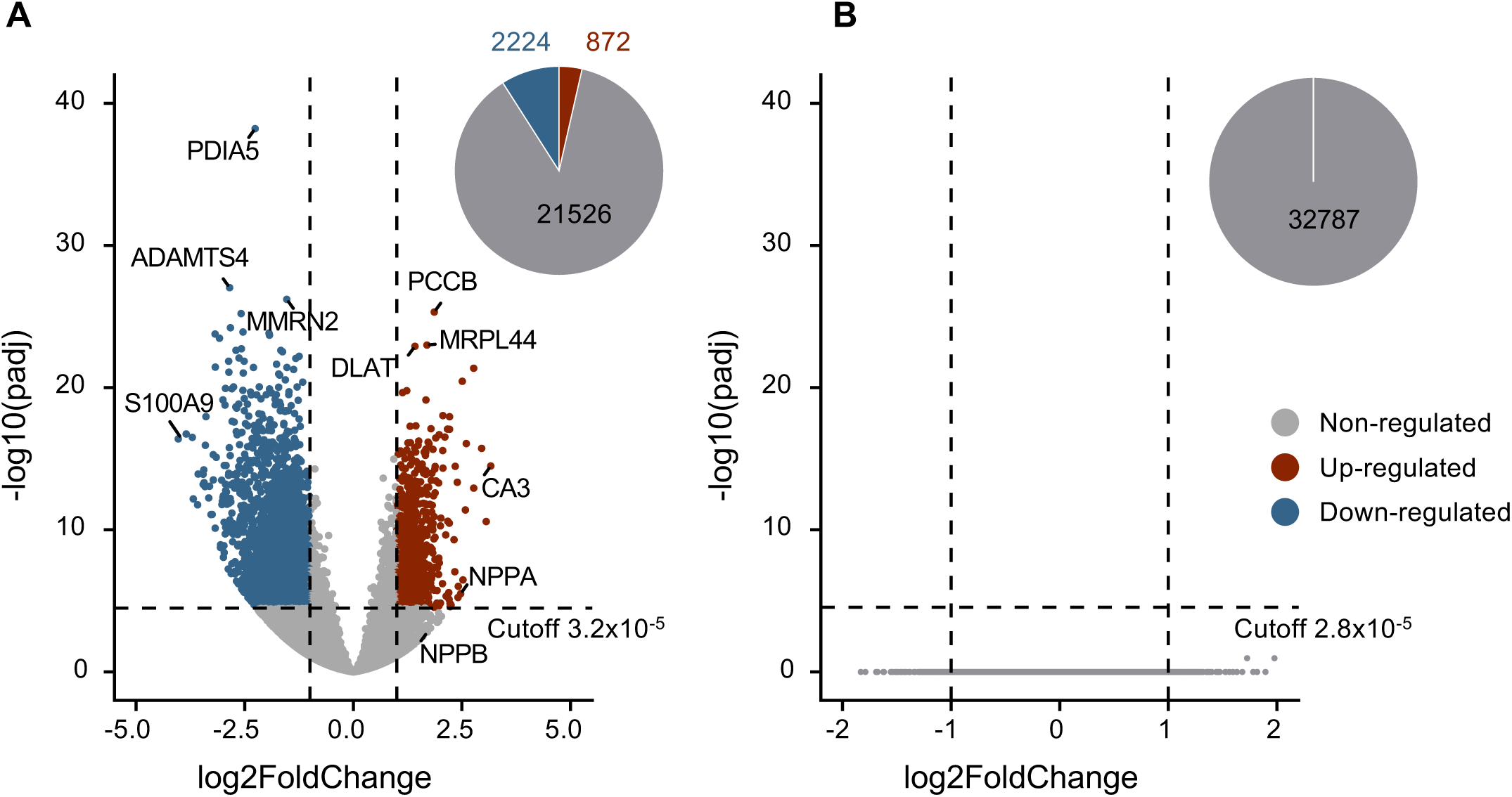
Differentially expressed genes. **A** Volcano plots of heart failure (HF) vs. control (CTRL) and **B** dilated cardiomyopathy (DCM) vs. ischemic cardiomyopathy (ICM). Significantly up-regulated, down-regulated and non-regulated genes are depicted in red, blue and grey, respectively. Vertical dashed lines mark the log fold-change cutoff of −1 and +1 and the horizontally dashed line shows the significance cutoff, with p-values calculated by p-value permutation. The number of genes in the different groups are displayed in the pie chart.

### Long-read sequencing uncovers thousands of novel splicing events

In total, 78,520 isoforms were detected with long-read sequencing and, out of those, 31% (n = 24,446) were novel and 69% (n = 54,074) were known (**Fig. 2A**). The GENCODE transcriptome was used as a reference to determine the distribution of known and novel isoforms. The length is displayed on a log-scale, revealing a prominent peak in the reference at approximately 2.8 kbp. The density of the known isoforms showed almost the same distribution, except for an additional larger peak around 3.4 kbp. However, novel isoforms tend to have the highest density for longer isoforms, which aligns with expectations based on the sequencing technology used (**Fig. 2B**). In our patient cohorts, the majority of genes had only one isoform annotated, however genes with more than one isoform were frequently found (**Fig. 2C**). Most of the detected transcripts are multi-exonic, indicating no bias towards smaller mono-exonic fragments (**Fig. 2D**). In addition, we determined the number of the most common alternative splicing events, including skipped exons (SE), alternative first (AF) and last (AL), as well as mutually exclusive (ME) exons and alternative 5’ and 3’ ends (A5 and A3, respectively). The splicing pattern was similar in novel compared to known isoforms with highest known counts at SE (novel=3,269; known=11,300) and AF (novel=2,011; known=9,005). While we also detected more A5 in known isoforms (novel=2,714; known=4,255); for all the other events, the numbers are higher for the novel isoforms compared to the known isoforms: A3 (novel=4,219; known=1,553); AL (novel=2,542; known=2542); MX (novel=903; known=542); RI (novel=2,402; known=825) (**Fig. 2E**).

**Figure 2:**
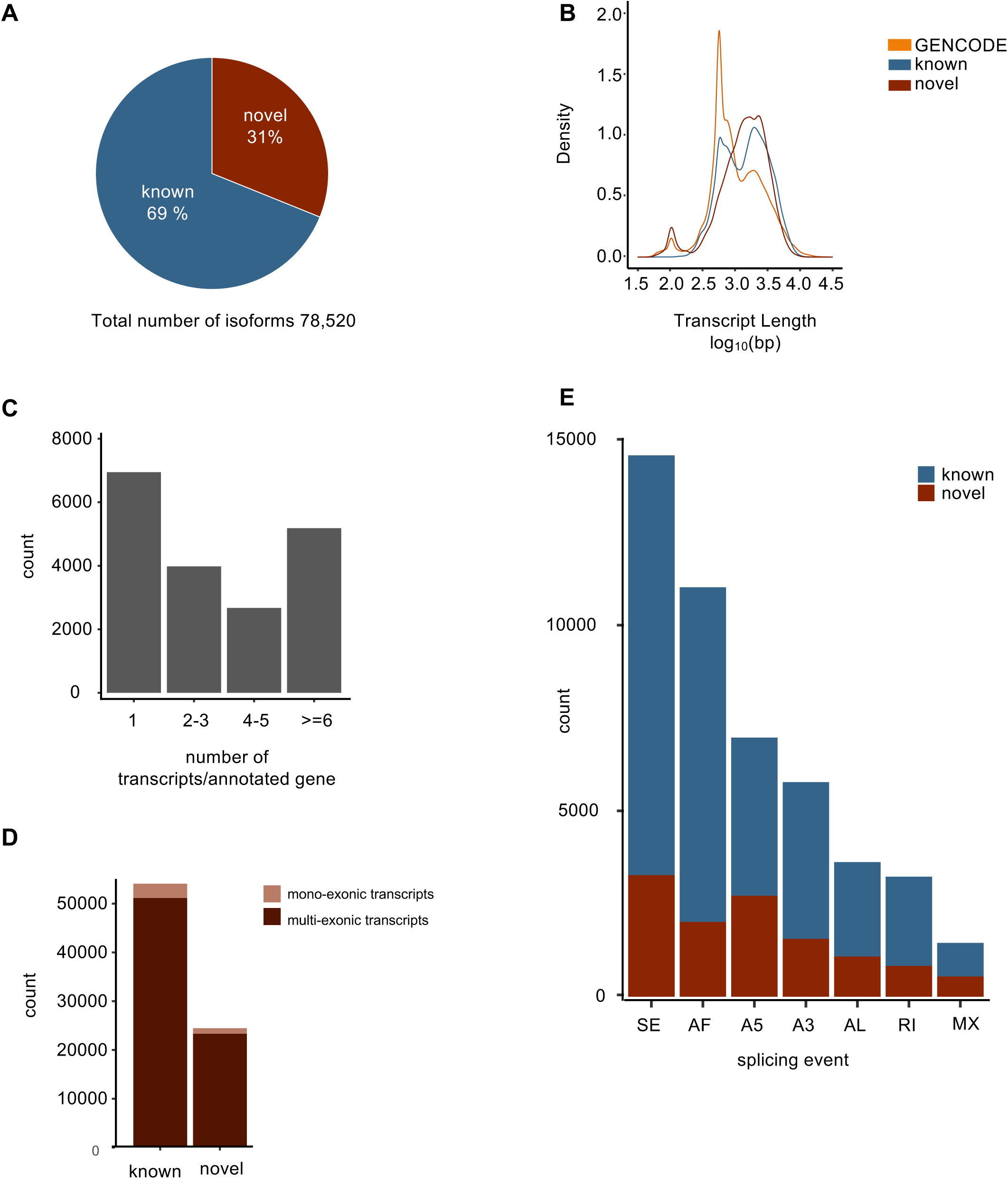
Long-read sequencing uncovers thousands of novel splicing events. **A** Total amount of transcripts identified by long-read sequencing, which detects 31 % as novel transcripts. **B** Kernal density read-length distribution of reference GENCODE transcripts (orange), known transcripts (blue), and novel transcripts (red). **C** Total counts of transcripts per annotated gene. **D** Total count of mono- and multiexonic transcript structure in known and novel transcripts. **E** Total count of splicing events as defined by SUPPA. Skipped Exon: SE, alternative first exon: AF, alternative 5’ end: A5, alternative 3’ end: A3, alternative last exon: AL, retained intron: RI, mutually exclusive exons: MX.

Next, we used SQANTi3 (Tardaguila *et al*, 2018) to classify the long-read transcripts where in total 44,132 transcripts had a full-splice match (FSM), of which 78.5% are coding (**Fig. 3A**). Additionally, we investigated the length distribution (median length 1,634 bp) which once again confirmed that there is no significant bias towards shorter fragments in the known isoforms with FSM compared to the novel ones in the catalog (NIC; median length 1,862 bp). The latter contains new combinations of already annotated splice junctions or novel splice junctions formed from already annotated donors and acceptors. Furthermore, a third group was identified, designated ‘novel not in catalog’ (NNC; median length 1,341 bp), which are transcripts that use novel donors and/or acceptors (**Fig. 3B**). The median length for the transcripts with incomplete splice-match (ISM) was slightly shorter (1,127 bp) due to the truncated reads. For the genic and fusion isoform groups (1,193.5 bp and 1,491 bp respectively), the median length is similar to known and novel isoforms in catalogue/not in catalogue. The antisense (158 bp), genic intron (70 bp), and the intergenic region (107 bp) have very short median read lengths and have a higher possibility of noise (**Fig. 3B**).

**Figure 3:**
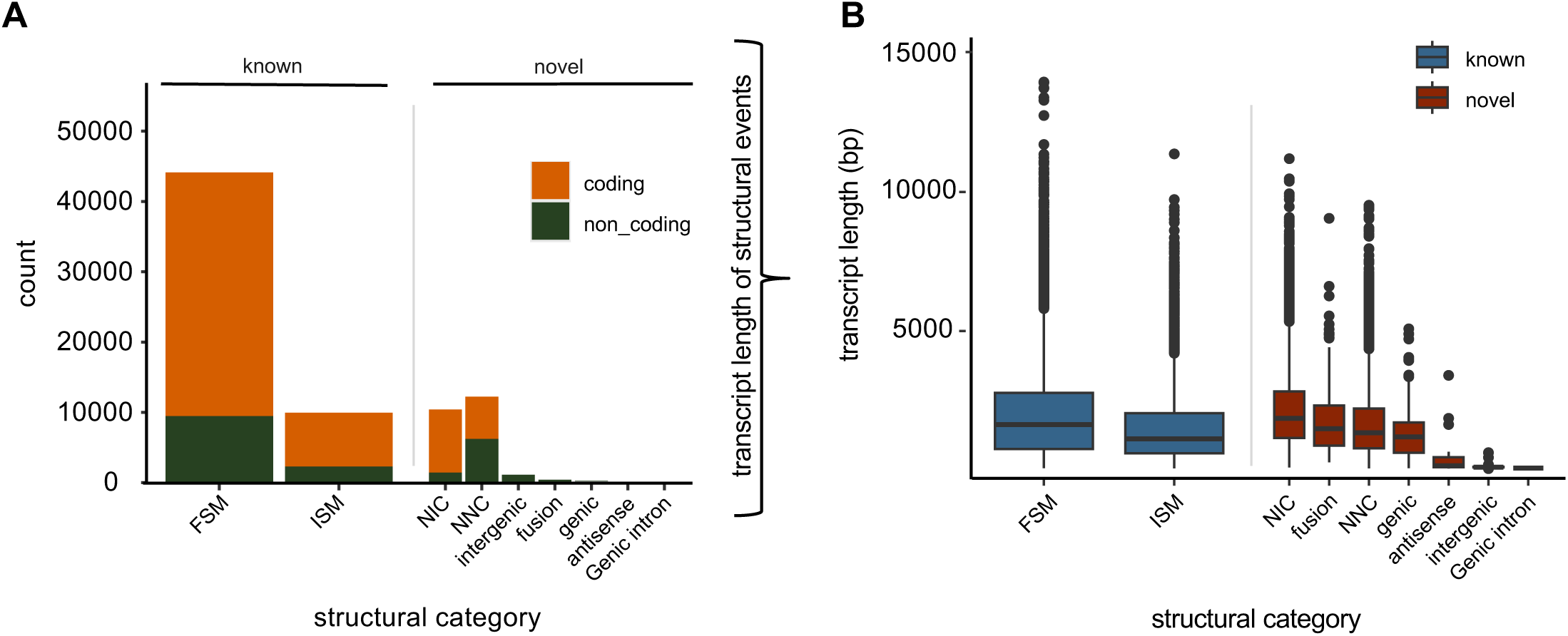
SQANTI classification of detected isoforms. **A** Total count (left) of structural events and **B** their length distribution (right) as defined by SQANTI. Known and novel isoforms are separated in coding (orange) and non-coding (green) isoforms. The known (blue) and novel (red) isoforms in the single structural categories were further analysed by their length. Full splice match: FSM, incomplete splice match: ISM, novel in catalog: NIC, novel not in catalog: NNC.

### Differentially expressed transcripts in human heart tissue

During heart failure, a shift occurs in myosin heavy chain (MHC) isoform expression, characterized by a decrease in α-MyHC and an increase in β-MyHC (Nakao *et al*, 1997). However, the detailed composition of the human sarcomere and the expression profiles of individual, mainly long isoform-encoding transcripts remain largely uncharacterized. Hence, we analysed the expression and isoform landscape of important sarcomeric genes in control and heart failure individuals. Based on the longread data, the significance cutoff was calculated by p-value permutation (padj= 2.0×10^−5^). Using this stringent cutoff, we identified 2,468 up-regulated and 4,505 down-regulated transcripts in DCM vs. CTRL (**Fig 4A**) and 1,749 up-regulated and 3,592 down-regulated transcripts in ICM vs. CTRL, (p-value= 2×10^−5^) (**Fig 4B**). Strikingly, no significantly differentially expressed transcript was detected between the two end-stage heart failure groups, again underlining that heart failure *per se* drives the transcriptional changes and not individual etiologies. In figures 4C and 4D, two examples of the isoform analysis in sarcomeric genes are displayed. In the cardiac *α* -actin gene (*ACTC1*), a novel 5’ end was detected and, five isoforms were differentially expressed in disease compared to control (**Fig 4C**). For the myosin-binding protein C3 gene (*MYBPC3*), a novel coding isoform was identified, which is formed by intron retention and is significantly down-regulated (**Fig. 4D**). As illustrated in the scheme in **Fig. 4E**, the majority of analyzed sarcomere transcripts exhibit significant changes.

**Figure 4:**
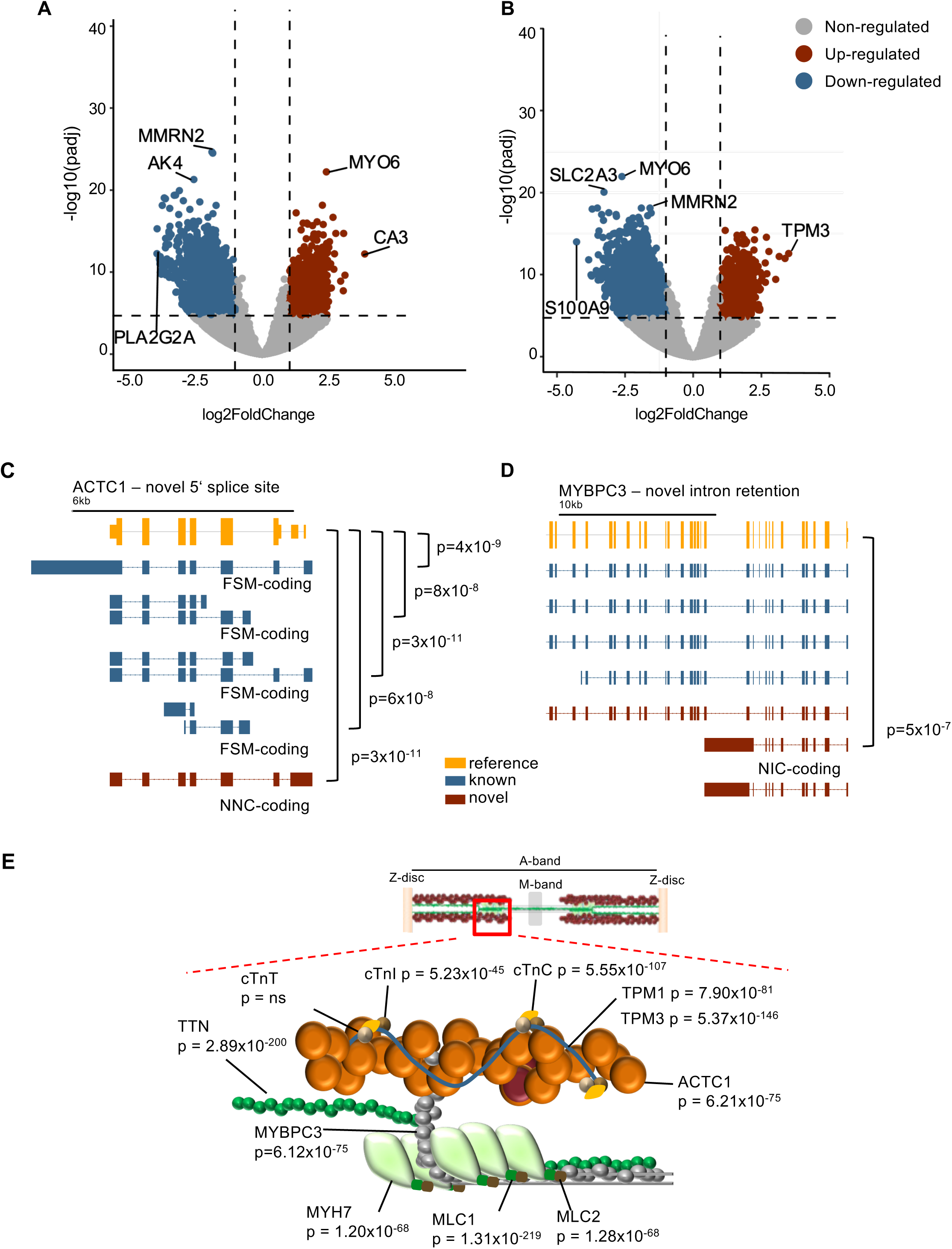
Differential transcript expression in heart failure. **A** Volcano plot of differentially expressed transcripts in DCM vs. CTRL and **B** volcano plots of differentially expressed transcripts in ICM vs. CTRL. Significantly up-regulated, down-regulated and non-regulated genes are depicted in red, blue and grey, respectively. Vertically dashed lines mark the log-fold change cutoff of −1 and +1 and the horizontally dashed line shows the significance cutoff, with p-values calculated by p-value permutation. The number of genes in the different groups is displayed in the pie chart. **C** Schematic transcript structure example of ACTC1. **D** Schematic transcript structure example of MYPC3. Reference gene, known isoform and novel isoform are shown in orange, blue and red, respectively. The p-values indicate the significant differentially regulated isoforms of heart failure vs. CTRL. FSM, NNC and NIC refer to SQANTI’s structural analysis and is only shown for the differentially expressed transcripts. **E** Scheme displaying that the most studied sarcomere transcripts shows significant changes in HF vs. CTRL. Sarcomere graphics were adapted from (Sedaghat-Hamedani *et al*, 2018).

### Tropomyosin 3 transcript usage is significantly different in heart failure vs. controls

Tropomyosins have been demonstrated to be integral to cardiomyocyte contractility and are of particular importance in mediating calcium-contraction coupling (Pieples *et al*, 2002). The four major tropomyosin genes—*TPM1*, *TPM2*, *TPM3*, and *TPM4*— produce a diverse array of isoforms through alternative splicing. The splicing of four key exons—1, 2, 6, and 9—varies across human tropomyosin isoforms and can be systematically represented using a four-letter code, where each letter corresponds to a specific splice variant of the respective exon. For example, “a.a.b.d” signifies that exon 1 is the splice form a, exon 2 is the splice form a, exon 6 is the splice form b, and exon 9 is the splice form d. If an exon is absent, it is represented by a dash (-), indicating a shorter isoform. For instance, “b.-.b.d” denotes the omission of exon 2, resulting in a truncated Tpm variant. Historically, the isoforms *TPM1*, *TPM2*, and *TPM3* primarily linked to muscle functions were referred to as α-Tpm, β-Tpm, and γ-Tpm, respectively (Wieczorek, 2018), while *TPM4* has been primarily linked to non-muscle functions. Notably, isoforms of *TPM1* and *TPM3* are also found in non-muscle cells, where they contribute to cytoskeletal organization and intracellular dynamics. While these genes have been extensively studied in animal models and during development, their full isoform diversity and specific roles in human tissues remain an area of ongoing research (Reindl *et al*., 2022; Schevzov G, 2011). While mutations in the *TPM3* are linked to congenital myopathies, the regulation of its major isoforms and their splice variants during heart disease remains poorly understood (Ohlsson *et al*, 2009). Our analysis revealed a significant prevalence of isoform switching in the tropomyosin genes in subjects diagnosed with heart failure, when juxtaposed with the control group. Under physiological conditions, *TPM1* is expressed at higher levels than *TPM3*, and during HF, the *TPM1-207* transcript encoding the long isoform Tpm1.1 (a.b.b.a) is specifically up-regulated (**Fig. 5A**). Intriguingly, *TPM3-224* displays much higher expression levels in HF, almost reaching levels of *TPM1* (**Fig. 5A**), resulting in an even higher fold-change of gene expression in HF (**Fig. 5B**). A comparison of the transcripts *TPM3-224,* which encodes the long isoform Tpm3.12 (a.b.b.a), and *TPM1-207* reveals that they are paralougs with highly conserved intron-exon structures (**Fig. 5C**). Both genes are also highly conserved on the protein level (97%), as shown by protein alignment (**Fig. 5D**). An analysis of the variable amino acids revealed that Tpm3.12 lacks putative or proven Serine/Threonine phosphorylation residues at seven locations, which might circumvent posttranslational regulatiory mechanisms.

**Figure 5:**
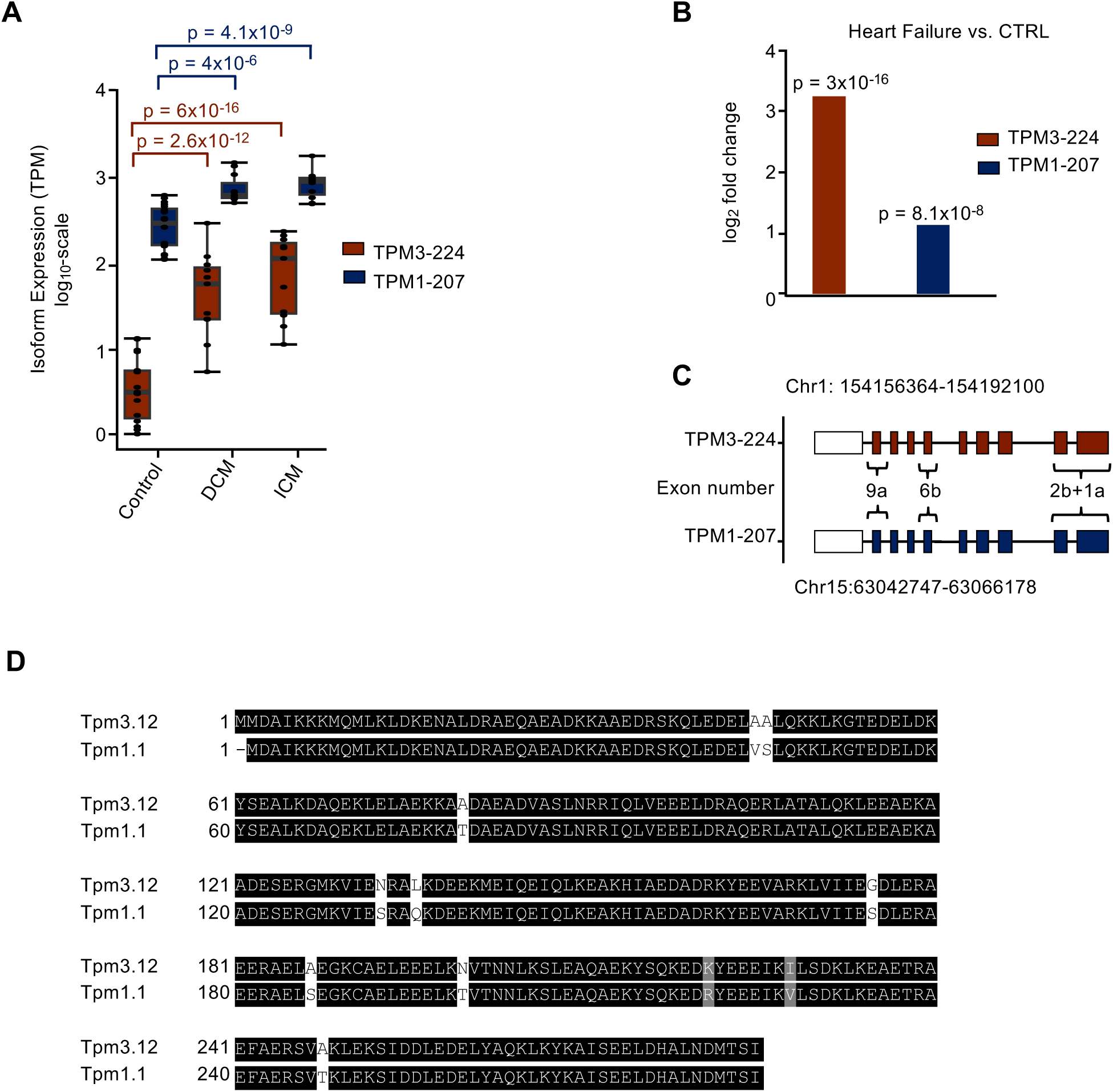
TPM3-224 is more up-regulated in heart failure compared to TPM1-1. 207. **A** Isoform expression in transcript per million (TPM) of TPM3-224 (red) and TPM1-207 (blue) in CTRL, DCM and ICM. Significance is calculated by Mann-Whitney U test. p-values are adjusted according to the Benjamini-Hochberg procedure and significance was accepted as p ≤ 0.05. **B** Significant log_2_ fold change of TPM3-224 (red) and TPM1-207 (blue) in heart failure (DCM + ICM) compared to controls as determined by DESeq2. p-values are Benjamini-Hochberg adjusted and the p-value cutoff = 2.3×10 was calculated by p-value permutation. **C** Highly similar isoform structure of TPM1-207 and TPM3-224 with nomenclature of important exons. Blue and red are in accordance with previously used colors for TPM1-207 and TPM3-224, respectively. White square indicates the UTR. **D** Protein sequence alignment of Tpm3.12 compared to Tpm1.1. Black squares indicate sequence similarity, whereas white squares indicate different amino acids and grey indicate amino acid belonging to the same chemical group.

We next analysed the precise composition of the isoforms of the HF-related *TPM3* upregulation. In healthy controls, *TPM3-206* (encoding the short isoform Tpm3.2 (b.-.b.d) and *TPM3-212* (encoding the short isoform Tpm3.1 (b.-.a.d)) are the predominantly produced transcripts. However, in the disease state, a notable shift in isoform expression occurs, with increased production of *TPM3-210*, which encodes Tpm3.13 (a.b.a.d), and *TPM3-224*, which encodes Tpm3.12 (a.b.b.a) (**Fig 6A**). In addition, transcript abundance in transcripts per million (TPM) was assessed for the four transcripts in CTRL, DCM, and ICM samples. In accordance with the differential gene expression analysis based on the negative binomial distribution (DESeq), we detected significant differences in HF vs. CTRL for *TPM3-224*: log_2_ fold change = 3.3, p-value = 1.5×10^−18^ and *TPM3-210*: log2 fold change = 3.1, p-value = 1.8×10^−17^. In DCM and ICM vs. CTRL, a significant up-regulation was detected for *TPM3-224*: log_2_ fold change = 3.03, p-value = 2.5×10^−12^ (DCM) and *TPM3-224*: log_2_ fold change = 3.5, p-value = 6×10^−16^ (ICM) and *TPM3-210*: log2 fold change = 2.9, p-value = 1.8×10^−11^ (DCM) and *TPM3-210*: log2 fold change = 3.4, p-value = 3.3×10^−15^ (ICM) was shown (**Fig 6B**). The differences between these transcripts arise from variations in exon 1, exon 6, and exon 9 (**Fig 6C**). According to official protein nomenclature, Tpm3.2 and Tpm3.1, the products of TPM3-206 and TPM3-212, respectively, are both classified as cytoplasmic tropomyosin isoforms. In contrast, Tpm3.12, encoded by TPM3-224, has been detected in striated muscle, whereas Tpm3.13, the product of TPM3-210, was previously reported only in rat and mouse but is now newly identified in human tissue (Geeves *et al*, 2015).

**Figure 6:**
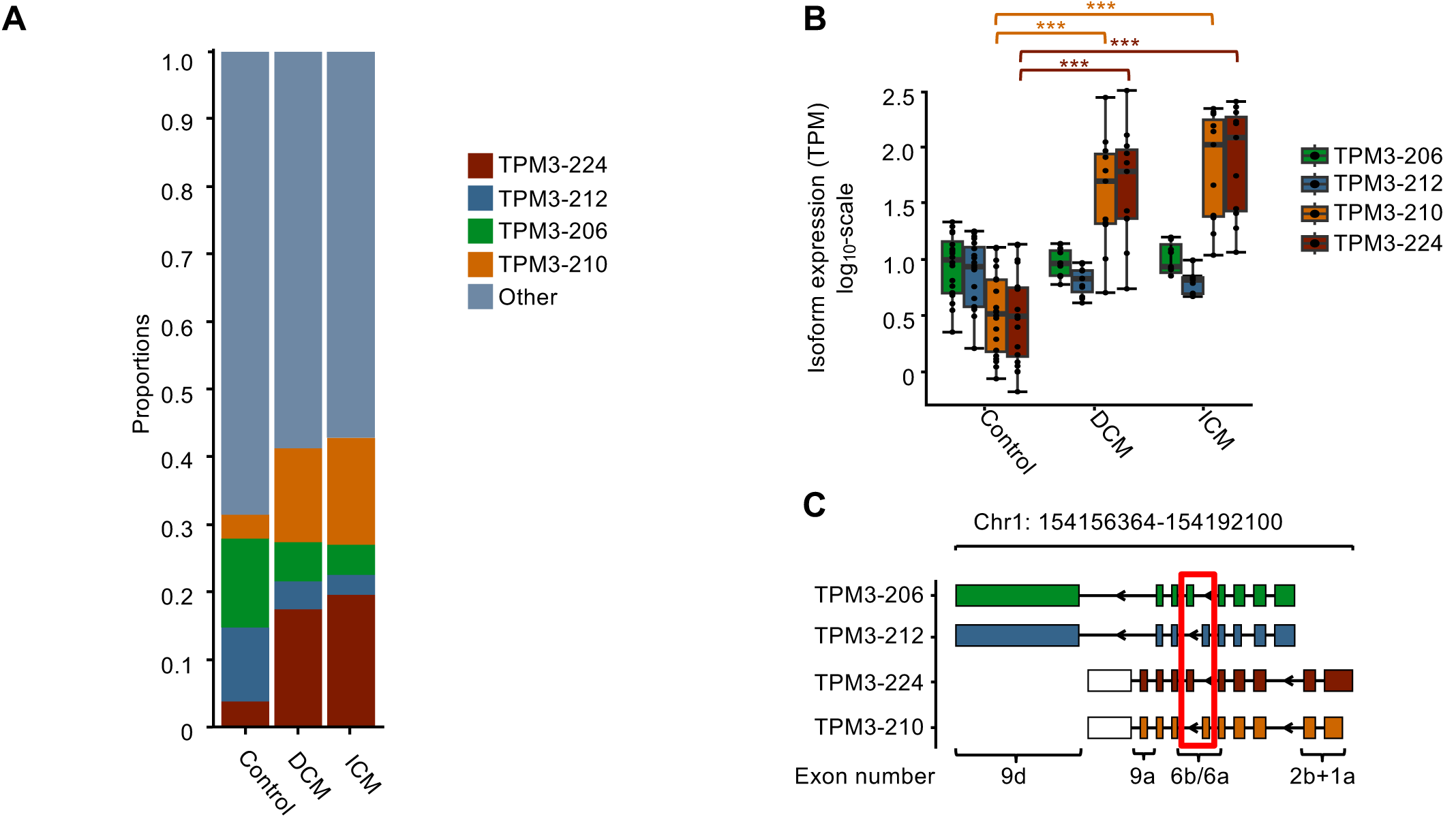
TPM3-224 is up-regulated in heart failure. **A** Proportions of *TPM3* transcripts, calculated using DRIMSeq. **B** Isoform expression levels measured in transcript per million (TPM). Statistical significance was determined by using the Mann-Whitney U test, with significance levels set as follows: p ≤0.001 = ∗∗∗. **C** Isoform structures highlighting the nomenclature of key exons. Red squares indicate exon 6.

### Tpm3.12 shows enhanced calcium responsiveness compared to Tpm1.1

The regulatory effect of tropomyosins on actin–myosin interaction can be investigated using *in vitro* motility assays, which measure the parameters such as the sliding velocity of actin and actin–tropomyosin filaments, driven by myosin motors (Kopylova *et al*, 2013; Kron & Spudich, 1986). Therefore, we established and validated motility assays to examine the effects of Tpm1.1 and Tpm3.12 under varying calcium conditions. All *in vitro*–synthesized proteins were separated on SDS-PAGE according to their molecular weights (**Fig 7A**). Using the motility assay, the v_max_ for Tpm1.1 was calculated to be 780.1 nm/s and the pCa_50_ was 6.3. For Tpm3.12, v_max_ was 728.5 nm/s and the pCa_50_ was 6.6 ( (**Fig 7B**).

**Figure 7:**
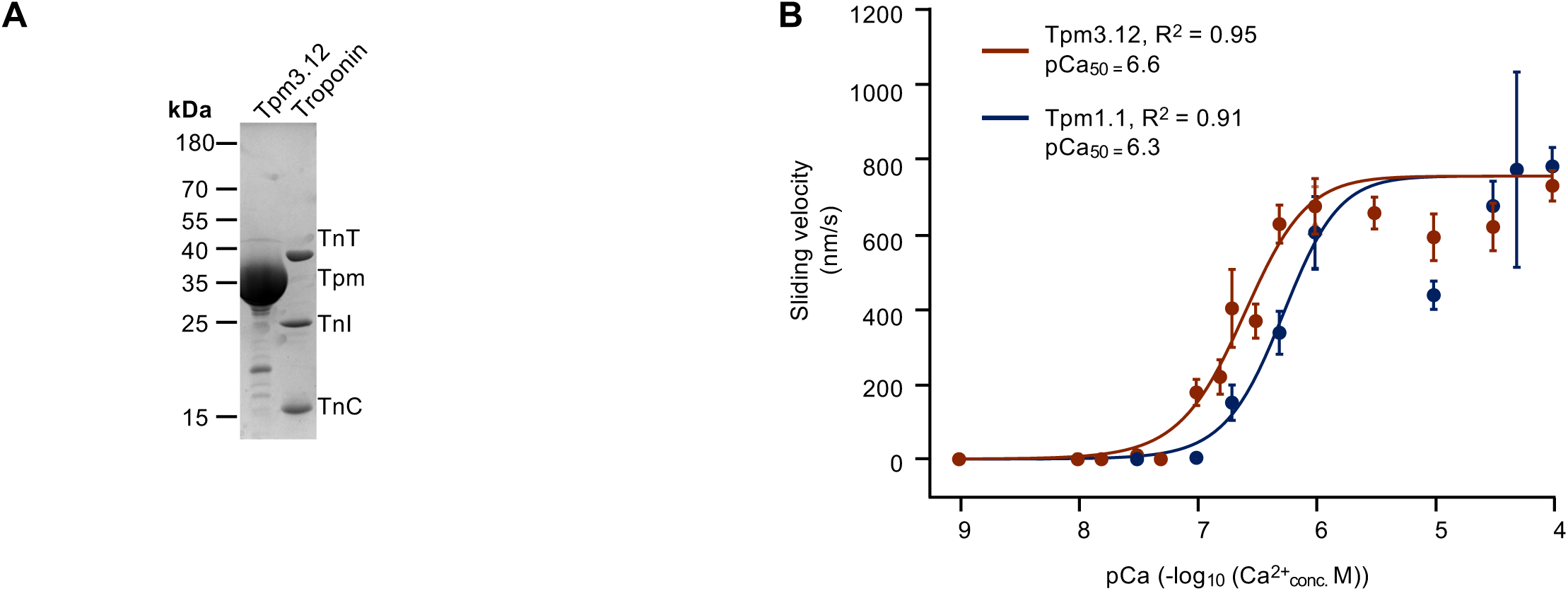
In-vitro motility assays performed in the presence of Tpm1.1 and Tpm3.12. **A** Proteins used for *in vitro* motility assays. SDS-PAGE of purified and reconstituted proteins are depicted. Expected molecular weights: Tpm3.12 (32.9 kDa), Troponin T (TnT, 34.5 kDa), Troponin C (TnC, 18.4 kDa), Troponin I (TnI, 24 kDa), Tpm1.1 (32.7 kDa), heavy meromyosin (HMM, 165 kDa), essential light chain (ELC, 22 kDa) and regulatory light chain (RLC, 16 kDa). PageRuler^TM^ (ThermFisher Scientific) was used as a molecular weight marker. **B** Calcium dose-response curve from *in vitro* motility assays for Tpm1.1 (blue) and Tpm3.12 (red). The coefficient of determination (R²) is provided for the Hill function fit. pCa values are expressed as - log_10_ of calcium concentrations (Molar) with pCa_50_ indicating the calcium concentration at half–maximal velocity. Sliding velocity values are presented as mean ±SEM.

## Discussion

Earlier investigations of the transcriptome reported distinct changes during heart failure (Sweet *et al*., 2018) (Zhu *et al*, 2022). While these studies relied on short-read sequencing with its high base-call accuracy (Shumate *et al*, 2022), correct mapping of isoforms is not precisely possible due to read lengths below 300 bp and large sequence homology between isoforms. Nanopore long-read sequencing technology has emerged as a powerful and evolving technique for the analysis of such isoform-specific transcriptome, enlightening alternative splicing and abundance of novel transcripts.

The regulation of cardiac transcript isoforms is governed by a complex interplay of splicing factors, cofactors, and upstream signaling pathways, which are essential for maintaining cardiac function. Central to this process are key splicing regulators, such as RBM20, SLM2, hnRNP A1, and RBFox1, whose interactions shape the expression of critical sarcomeric proteins, particularly the tropomyosin isoforms (**Table 1**). RBM20, for instance, represses *TPM3* gene products by recruiting U2 snRNP and PTBP1, thus favoring the expression of the cardiac-specific *TPM1-207* transcript. This regulation of tropomyosin isoforms is pivotal for maintaining proper sarcomere function in the heart. In contrast, SLM2 and hnRNP A1 promote the expression of non-muscle *TPM3* gene products by disrupting U2AF65 recruitment, highlighting their role in altering the balance between muscle and non-muscle isoforms in response to various physiological and pathological conditions. Furthermore, RBFox1 enhances *TPM1-207* expression by stabilizing the U1 snRNP complex and binding to UGCAUG enhancer motifs in sarcomeric transcripts, further ensuring the maintenance of sarcomere integrity.

**Table 1.** Splicing Machinery Regulating Tropomyosin Isoforms in Human Left Ventricle Cells.

| Isoform | Transcript | Primary Splicing Regulator | Supporting Regulators | Splicing Complexes & Cofactors | Upstream Signaling Pathways | Disease Associations |
| --- | --- | --- | --- | --- | --- | --- |
| **Tpm1.1** | *TPM1-207* | RBM20 | MBNL1, SRSF1, SRSF2, RBFOX1 | RBM20-repressed spliceosome, U2 snRNP, PTBP1 | PKC, ERK/MAPK, RBM20-mediated exon repression | Heart failure, hypertrophic cardiomyopathy (HCM), dilated cardiomyopathy (DCM) |
| **Tpm3.12** | *TPM3-224* | SLM2 | hnRNP A1, SRSF1, CELF1 | hnRNP-enriched splicing complexes, CELF1-dependent spliceosome | SLM2-driven exon inclusion, ERK/MAPK, Calcium signaling | Dilated cardiomyopathy (DCM), cytoskeletal instability |
| **Tpm3.13** | *TPM3-210* | SLM2 | MBNL1, hnRNP A1 | MBNL1-associated spliceosome, U2AF65 | SLM2-dependent exon switching, TGF-β signaling | Cardiac fibrosis, myofibril disorganization |
| **Tpm3.1** | *TPM3-212* | RBFOX1 | MBNL1, MBNL2 | RBFOX1-bound U1 snRNP, UGCAUG-dependent enhancer complex | RhoA/ROCK, Actin cytoskeletal regulation | Vascular remodeling post-MI, altered fibroblast function |
| **Tpm3.2** | *TPM3-206* | SLM2 | hnRNP A1, SRSF1 | hnRNP A1/U2AF65-mediated splicing machinery | SLM2-mediated exon selection, TGF-β signaling | Fibrotic heart disease, increased ECM deposition |

Upstream signaling pathways, including those mediated by PKC, ERK/MAPK, and calcium signaling, also play a critical role in regulating the splicing of TPM isoforms. RBM20 is known to be regulated by both PKC and ERK/MAPK signaling, ensuring that *TPM1-207* remains the dominant isoform in the left ventricle of a normally functioning heart. The activation of SLM2 by ERK/MAPK and calcium signaling pathways leads to an increase in TPM3 isoform expression, which is particularly important in the context of cardiac stress. Additionally, TGF-β signaling and the RhoA/ROCK pathway regulate TPM3 isoforms, especially during fibrosis and post-injury remodeling, underscoring the dynamic nature of isoform expression during heart disease.

The relevance of these splicing mechanisms extends beyond basic biology, with mutations and dysregulation of these factors contributing to cardiac disease. Mutations in RBM20, for example, lead to a shift from *TPM1-207* to *TPM3* transcripts, weakening sarcomere function and contributing to dilated cardiomyopathy (DCM) and heart failure. Similarly, overexpression of SLM2 in heart failure results in increased levels of *TPM3* gene products, driving cytoskeletal remodeling and fibrosis. In cases of cardiac hypertrophy, downregulation of RBFox1 results in the loss of *TPM1-207*, leading to compromised sarcomere integrity and activation of fibroblasts, further contributing to the progression of heart disease.

Recent advances in transcriptome analysis, particularly through the use of long-read sequencing technologies, have provided deeper insights into the complexity of cardiac splicing regulation. Previous studies on RBM20 mutant hiPS-CM, analyzed with FulQuant (Zhu et al., 2021), and the pig cardiac transcriptome, examined with StringTie (Müller et al., 2021), have identified thousands of isoforms previously unknown. In our own study, we analyzed the cardiac transcriptomes of more than 30 human individuals, providing the most detailed and unbiased view of the human isoform landscape in the heart. Our stringent cut-offs, designed to reduce the risk of false positives, combined with the use of FLAMES after careful benchmarking of available tools, ensure the reliability of our findings. These protocols not only enhance our understanding of the human cardiac isoform landscape but also pave the way for the application of nanopore sequencing in human diagnostic testing (Sedaghat-Hamedani et al., 2022).

As demonstrated in transcript redundancy analysis, *PCCB, MRPL44, DLAT, CA3* and *NPPA* were found to be among the most significantly up-regulated genes in HF. Propionyl-CoA carboxylase (PCC) is a dodecameric enzyme complex, consisting of 6α- and 6β-subunits, which are encoded by the *PCCA* and *PCCB* genes. The deficiency of PCC, which is encoded by *PCCB*, leads to propionic acidemia resulting in DCM (Riemersma *et al*, 2017). The protein product of the *MRPL44* gene, ml44, was identified as a component of the large subunit of the mitochondrial ribosome (mitoribosome) and pathogenic variants in *MRPL44* causes infantile cardiomyopathy due to a mitochondrial translation defect (Friederich *et al*, 2021). Carbonic anhydrase (CA) is a zinc-containing enzyme that catalyzes the reversible hydration of carbon dioxide and that plays a role in regulating the transport of bicarbonate (Lindskog, 1997). In a recent study, CA2 and CA3 were found to be more highly expressed in the HF group compared to controls, as determined by ELISA (Su *et al*, 2021). Carbonic Anhydrase 3 (CA3) was shown to be required for cardiac repair post-myocardial infarction via Smad7-Smad2/3 signaling pathway (Su *et al*, 2024). In this study, the authors observed that CA3 deficiency restrains collagen synthesis, cell migration and gel contraction of cardiac fibroblasts, whereas overexpression of CA3 improves wound healing and cardiac fibroblast activation. Accordingly, the detected upregulation of CA3 in our DCM patients might point to an activation of a repair mechanism. Dihydrolipoamide S-Acetyltransferase (*DLAT)* encodes the component E2 of the multi-enzyme pyruvate dehydrogenase complex (PDC). PDC resides in the inner mitochondrial membrane and catalyzes the conversion of pyruvate to acetyl coenzyme A. Interestingly, we have previously identified altered pyruvate metabolite levels in an ambulatory HF cohort (Haas *et al*, 2021). Both *NPPA* and *NPPB* are induced by cardiac stress and serve as markers for cardiovascular dysfunction or injury (Giovou *et al*, 2024). In our analysis, the *NPPA* gene was found to be significantly up-regulated while the *NPPB* gene remained unaltered.

Among the strongly down-regulated genes, *ADAMTS4* and *S100A9* were both reported in previous studies as cardiac injury markers. The disintegrin-like and metalloproteinase with thrombospondin motif (ADAMTS) family comprises 19 proteases that regulate the structure and function of extracellular proteins in the extracellular matrix and blood. A number of studies have also investigated the potential role of ADAMTS-1, −4 and −5 in cardiovascular disease (Santamaria & de Groot, 2020). *ADAMTS4* was identified as a novel adult cardiac injury biomarker with therapeutic implications in patients with cardiac injuries. It is also downregulated, as we observed in our study, and depicted among the hub genes identified in potential DCM-related targets by meta-analysis and co-expression analysis of human RNA-sequencing datasets (Yuan et al, 2022).

During HF, the isoform switch of myosin heavy chains (MHC) plays a crucial role in the alteration of cardiac contractility. In a healthy heart, the predominant MHC isoform is alpha-myosin heavy chain (α-MHC), which supports efficient contraction due to its rapid ATPase activity. However, in heart failure, a shift occurs towards the expression of the beta-myosin heavy chain (β-MHC), a slower isoform associated with reduced contractile function (Liu *et al*, 2016; van der Velden *et al*, 2003). This isoform transition is believed to be a compensatory response to maintain cardiac function under stress, but it leads to a decline in the heart’s efficiency and performance. The molecular mechanisms underlying this isoform switch involve changes in the transcriptional regulation of the MHC genes, including alterations in the expression of factors such as myocyte enhancer factor 2 (MEF2) and the calcium/calmodulin-dependent kinase II (CaMKII) pathway, both of which were implicated in promoting β-MHC expression in the failing heart (Sato et al., 2003; Helm et al., 2014). We observed in this study a profound change of sarcomeric isoforms that might functionally impact cardiomyocytes in a similar manner. Alternative splicing of *TPM* genes has been previously reported to generate tropomyosin isoforms with distinct biochemical and functional properties (Moraczewska, 2020) (Kengyel *et al*, 2024; Pathan-Chhatbar *et al*, 2018; Rajan *et al*, 2010). Previous investigations suggested only limited expression of *TPM3* in the heart (Dube et al, 2020) (Tucholski et al, 2020) (Lawlor et al, 2010). By performing single-nucleus RNA sequencing of nearly 600,000 nuclei at a single cell resolution, *TPM3* expression was specifically detected in cardiomyocytes across samples from 11 DCM and 15 HCM patients; as well as 16 non-failing hearts (Chaffin et al, 2022). This observation suggested that *TPM3* is expressed in cardiomyocytes across both diseased and healthy heart tissues, potentially indicating a role in cardiac function or disease pathology. The analysis could, however, not incorporate nanopore long-read data. Hence, the present data underscores the identification of hitherto unrecognized tropomyosin isoforms. *TPM3* gene products are among the most highly responsive in heart failure, approaching expression levels comparable to the dominant *TPM1* gene products. In the present study, *in vitro* motility assays were conducted to investigate the calcium sensitivity of Tpm3.12 and Tpm1.1. The results of these assays reveal that Tpm3.12 exhibits greater calcium sensitivity compared to Tpm1.1.

Collectively, our study shows an orchestrated remodeling of the cardiac isoform landscape during heart failure. The changes are highly reproducible and comparable in ischemic and non-ischemic causes. With novel compounds being tested in clinical trials, it becomes more crucial to understand the targeted isoforms and their variations under different disease conditions. This might improve efficiency and reduce unwanted side effects. Especially the cardiac sarcomere seems highly flexible regarding its isoform repertoir.

### Limitations

A limitation of this study is the use of bulk RNA sequencing, which captures transcriptomic changes from whole tissue rather than individual cardiomyocytes. While non-myocytes contribute to heart tissue remodeling in heart failure, their influence on transcript abundance is likely limited due to the dominant transcriptional output of cardiomyocytes (Litviňuková et al., 2020). Single-cell and single-nucleus RNA sequencing studies have demonstrated that key sarcomeric transcript changes, including isoform switching in *TPM1* and *TPM3*, are largely preserved across methods, suggesting that the observed shifts primarily reflect cardiomyocyte remodeling (Tabula Sapiens Consortium, 2022; Koenig et al., 2022). However, bulk sequencing cannot fully resolve cell-type-specific contributions, and future studies using single-cell approaches will be essential to confirm the cardiomyocyte-specific nature of these isoform changes.(Cadosch *et al*, 2024; Koenig *et al*, 2022; Tabula Sapiens *et al*, 2022)

## Materials and Methods

### Human left ventricular heart tissue

The present study has been approved by the ethics committee of the Medical faculty of Heidelberg University (appl. no. S-390/2011) and participants have given written informed consent. Control samples were used according to the protected health information (45 C.F.R. 164.514 e2) (bioserve) and the BCI informed consent F-641-5 (biochain).

### RNA isolation

Heart pieces were stored at −80 °C and RNA was extracted from 200 ng tissue using the AllPrep DNA/RNA/miRNA Kit (Qiagen, #80204). RNA concentrations were determined on Agilent Technology’s Fragment Analyser with either the standard sensitivity (range 5 - 500 ng/μL) or the high sensitivity kit (range 50 −5000 pg/μL) according to manufacturer’s protocol. RNA quality number (RQN) was measured to assess the quality of the RNA.

### Generation of nanopore cDNA libraries and sequencing

For nanopore sequencing, 30 ng of mRNA was mixed with 0.3 ng spike-in control Lexogen SIRV E0 mix (#050.01). The libraries were prepared according to the manufacturer’s protocol for the DCS-109 kit, however, small changes were introduced. In brief, the first step of the library preparation involves cDNA synthesis with a polydT primer (VNP offered by Nanopore) and the maxima H reverse transcriptase (Thermo Scientific^TM^, #EP0751). Due to maxima Hs terminal transferase activity, it is possible to use the additional bases as an anchoring site for a strand switching primer (SSP, provided by Nanopore) to synthesize the opposite strand as well. Primers complementary to the polydT primer and SSP primer overhangs (provided by Nanopore) were used for the LongAmp® *Taq* Polymerase PCR. Followed by an exonuclease I (20 units, NEB, #M0293) digestion, PCR products were purified with AMPure XP beads (Beckman Coulter, #A63882). DNA repair to obtain 5’phosphorylated and 3’dA-tailed sites was performed with Ultra II End-prep module from NEB (#E7546S) and DNA was purified with 60 μL AMPure XP beads. The provided nanopore ONT adapters were ligated and 60 ng final library was prepared according to manufacturer’s instructions and loaded on a primed GridION flowcell (R9.4.1). Sequencing was performed with one sample per flowcell for 48 h on the GridION which basecalls directly with Guppy.

### Protein expression and purifications

Tropomyosins were overexpressed by transforming *E. coli* BL21(DE)-pNatB with tropomyosin constructs cloned in either the pET-23a (+) vectors or pETpRha vectors. The pNatB plasmid encodes the regulatory subunit naa25⁺ and the catalytic subunit naa20⁺ of the NatB complex from fission yeast, both under the control of a T7 promoter (Johnson *et al*, 2010). Actin was isolated from rabbit skeletal muscle and acetone powder was prepared according to (Lehrer & Kerwar, 1972). G-actin was extracted from the acetone powder and polymerized into F-actin by adding polymerization mix (2 M KCl and 40 mM MgCl_2_) to a final concentration of 100 mM KCl and 2 mM MgCl_2_. The mixture was incubated at room temperature for 1 hour before being stored on ice. Human troponin subunits (cTnC, cTnI, cTnT) were cloned into pET11cvectors containing ampicillin and chloramphenicol resistance and overexpressed in *E.coli* BL21 (RosettaDE3) pLysS cells. Purification of cTnI and cTnC was performed according to (al-Hillawi *et al*, 1994), while cTnT purification followed the protocol described by (Krüger *et al*, 2003).

### *In vitro* motility assay

Fluorescently labelled actin gliding assays on surface-bound *β* -cardiac myosin HMM were conducted as previously described (Reindl *et al*., 2022) using an Olympus IX70 fluorescent microscope. In brief, 0.2 mg/ml *β* -cardiac myosin HMM was diluted in assay buffer (25 mM MOPS, 50 mM KCl, 5 mM MgCl_2_, 10 mM DTT, pH 7.4) and bound to the cover slip of a flow cell pre-coated using 1% nitrocellulose dissolved in pentyl-acetate. To minimize nonspecific actin binding, the surface was blocked with 0.5 mg/ml bovine serum albumin. F-actin labeled with 20 nM Atto550-Phalloidin (Merck) was incubated on the surface for 3 minutes under rigor conditions. Motility was initiated by adding 2 mM ATP in an assay buffer supplemented with 0.5% methylcellulose and an anti-photobleaching mix (5 mg/ml glucose, 100 µg/ml glucose-oxidase, 100 µg/ml catalase). At least three video sequences were captured using Olympus xcellence RT software at either one or two frames per second. The velocity and filament length distribution of at least 300 actin filaments were analyzed using FiJi software with a customized version of the wrMTrck plugin (Nussbaum-Krammer *et al*, 2015; Schindelin *et al*, 2012). For the measurements involving Ca^2+^ ions, free Ca^2+^ ion concentrations were calculated using MAXCHELATOR with the following parameters: 37 °C, 0.055 N ion contribution, pH = 7.4, ATP = 0.002 M, EGTA = 0.001 M and 0.004 M Mg^2+^

## Acknowledgements

This study was supported by CRC 1550 (Molecular Circuits of Heart Disease) and the Leducq Foundation (CASTT grant). D.J.M was supported by grants from Deutsche Forschungsgemeinschaft (Project number 462266917) and the German Federal Ministry of Education and Research under Grant Agreement 01GM1922B.

## Author contributions

### Data availability

All data is avalaible upon request.

